# NeuroPycon: An open-source Python toolbox for fast multi-modal and reproducible brain connectivity pipelines

**DOI:** 10.1101/789842

**Authors:** David Meunier, Annalisa Pascarella, Dmitrii Altukhov, Mainak Jas, Etienne Combrisson, Tarek Lajnef, Daphné Bertrand-Dubois, Vanessa Hadid, Golnoush Alamian, Jordan Alves, Fanny Barlaam, Anne-Lise Saive, Arthur Dehgan, Karim Jerbi

**Author notes:** equal contribution (David Meunier, Annalisa Pascarella).

## Abstract

Recent years have witnessed a massive push towards reproducible research in neuroscience. Unfortunately, this endeavor is often challenged by the large diversity of tools used, project-specific custom code and the difficulty to track all user-defined parameters. NeuroPycon is an open-source multi-modal brain data analysis toolkit which provides Python-based template pipelines for advanced multi-processing of MEG, EEG, functional and anatomical MRI data, with a focus on connectivity and graph theoretical analyses. Importantly, it provides shareable parameter files to facilitate replication of all analysis steps. NeuroPycon is based on the NiPype framework which facilitates data analyses by wrapping many commonly-used neuroimaging software tools into a common Python environment. In other words, rather than being a brain imaging software with is own implementation of standard algorithms for brain signal processing, NeuroPycon seamlessly integrates existing packages (coded in python, Matlab or other languages) into a unified python framework. Importantly, thanks to the multi-threaded processing and computational efficiency afforded by NiPype, NeuroPycon provides an easy option for fast parallel processing, which critical when handling large sets of multi-dimensional brain data. Moreover, its flexible design allows users to easily configure analysis pipelines by connecting distinct nodes to each other. Each node can be a Python-wrapped module, a user-defined function or a well-established tool (e.g. MNE-Python for MEG analysis, Radatools for graph theoretical metrics, etc.). Last but not least, the ability to use NeuroPycon parameter files to fully describe any pipeline is an important feature for reproducibility, as they can be shared and used for easy replication by others. The current implementation of NeuroPycon contains two complementary packages: The first, called *ephypype,* includes pipelines for electrophysiology analysis and a command-line interface for on the fly pipeline creation. Current implementations allow for MEG/EEG data import, pre-processing and cleaning by automatic removal of ocular and cardiac artefacts, in addition to sensor or source-level connectivity analyses. The second package, called *graphpype,* is designed to investigate functional connectivity via a wide range of graph-theoretical metrics, including modular partitions. The present article describes the philosophy, architecture, and functionalities of the toolkit and provides illustrative examples through interactive notebooks. NeuroPycon is available for download via github (https://github.com/neuropycon) and the two principal packages are documented online (https://neuropycon.github.io/ephypype/index.html. and https://neuropycon.github.io/graphpype/index.html). Future developments include fusion of multi-modal data (eg. MEG and fMRI or intracranial EEG and fMRI). We hope that the release of NeuroPycon will attract many users and new contributors, and facilitate the efforts of our community towards open source tool sharing and development, as well as scientific reproducibility.

## 1. Introduction

Recent years have witnessed a massive push towards reproducible experiments in neuroscience. Some of the leading projects include OpenfMRI (now OpenNeuro, Poldrack & Gorgolewski, 2017), allowing researchers worldwide to test hypothesis on a massive cohort, NeuroSynth (Yarkoni et al., 2011) a brain mapping framework to automatically conduct large-scale, high-quality neuroimaging meta-analyses or the development of the Nipype framework (Gorgolewski et al., 2011), a very useful tool developed initially for the MRI field and which facilitates data analysis by wrapping commonly-used neuroimaging software into a common Python framework. Nipype was originally designed to provide rapid comparative development of algorithms and to reduce the learning curve necessary to use different packages. Nipype’s original intention was to provide a concrete tool to respond to some criticisms of the neuroscientific community as a whole for the lack of reproducibility of experiments, in particular when it comes to attempts to reproduce research on a wider scale both in basic research and clinical trials (Gilmore et al, 2017). One step forward in this direction is also to provide the source code used for a given analysis pipeline, not to mention sharing data in standard formats (Poline et al, 2012; Gorgolewski et al, 2015).

The release of Nipype was a major step forward allowing researchers to wrap virtually all fMRI and MRI software into a common framework, where all analysis steps and, critically, all the parameter settings are tractable and easily accessible. Nipype is for instance used to wrap, into a common pipeline, some of the most frequently used MRI research tools, ranging from SPM (Penny et al., 2011), FSL (Smith et al., 2004) and AFNI in functional MRI (Cox 1996), ANTS (Avants et al., 2009), Freesurfer in structural MRI (Dale et al., 1999), Camino (Cook et al., 2006) and Mrtrix in diffusion imaging (Tournier et al., 2012). The advantages of having a single unified framework are particularly obvious, especially when it comes to scaling up and/or reproducing neuroimaging research.

Interestingly, while encouraging progress has been achieved with such strategies in the fMRI field, similar initiatives for the fields of MEG and EEG data analysis are still in their early days (Bigdely-Shamlo et al., 2015; Andersen, 2018; Jas et al., 2018; Niso et al., 2019). Yet, certain MEG and EEG analyses consist of a sequence of processing steps that may involve several software packages and possibly multiple programming languages and environments. An MEG analysis that starts from raw data and ends up with group-level statistics may, for example, require the use of *Freesurfer* for cortical segmentation, *MNE* for preprocessing and source estimation, *radatools* for connectivity metrics and *Visbrain* (Combrisson et al., 2019) for 3D visualizations. Thus, most of the processing is conducted using multiple heterogeneous software and in-house or custom tools leading to workflows that are hard or even impossible to reproduce in practice. Furthermore, with the exponential increase in data dimensionality and complexity, conducting state-of-the-art brain network analyses using MEG and EEG is becoming an increasingly challenging and time-consuming endeavor. Streamlining all the steps of MEG/EEG data analyses into a unified, flexible and fast environment could greatly benefit large-scale MEG/EEG research and enhance reproducibility in this field.

Here, we describe NeuroPycon, a free and open-source Python package, which allows for efficient parallel processing of full MEG and EEG analysis pipelines which can integrate many available tools and custom functions into a single workflow. The proposed package uses the Nipype engine framework to develop shareable processing pipelines that keep track of all analyses steps and parameter settings. Although initially developed with MEG and EEG in mind, NeuroPycon allows for multi-modal brain data analysis thanks to the flexibility and modularity it inherits form Nipype.

This paper describes the architecture, philosophy and rationale behind NeuroPycon. The functionalities of *ephypype* and *graphpype* are illustrated by describing how NeuroPycon is used to wrap existing tools that analyse electrophysiological data (e.g. MNE-python, Gramfort et al., 2013) and that perform graph-theoretical analysis (radatools^1^). Of course, wrapping many other software tools or packages within NeuroPycon workflows is possible by writing new Nipype interfaces to the desired tool.

## 2. Pipeline design, data structure and analysis workflow

### 2.1 Overview

#### Description

NeuroPycon provides computationally efficient and reusable workflows for advanced MEG/EEG and multi-modal functional connectivity analysis pipelines. Because NeuroPycon is powered by the Nipype engine (Gorgolewski, et al, 2015) it benefits from many of its strengths and shares the same philosophy. The NeuroPycon workflows expand and promote the use of the Nipype framework to the MEG/EEG research community. NeuroPycon links different software packages through nodes connected in acyclic graphs. (Fig 1A) The output of one pipeline can be provided as inputs to another pipeline. Furthermore, NeuroPycon is designed to process subjects in parallel on many cores or machines; If the processing is interrupted due to an error, NeuroPycon will only recompute the nodes which do not have a cache (Fig 1B). The current release of the NeuroPycon project includes two distinct packages: The *ephypype* package includes pipelines for electrophysiological data analysis and a command-line interface for on the-fly pipeline creation. A second one called *graphpype* is designed to investigate functional connectivity via a wide range of graph-theoretical metrics, including modular partitions.

**Figure 1.**
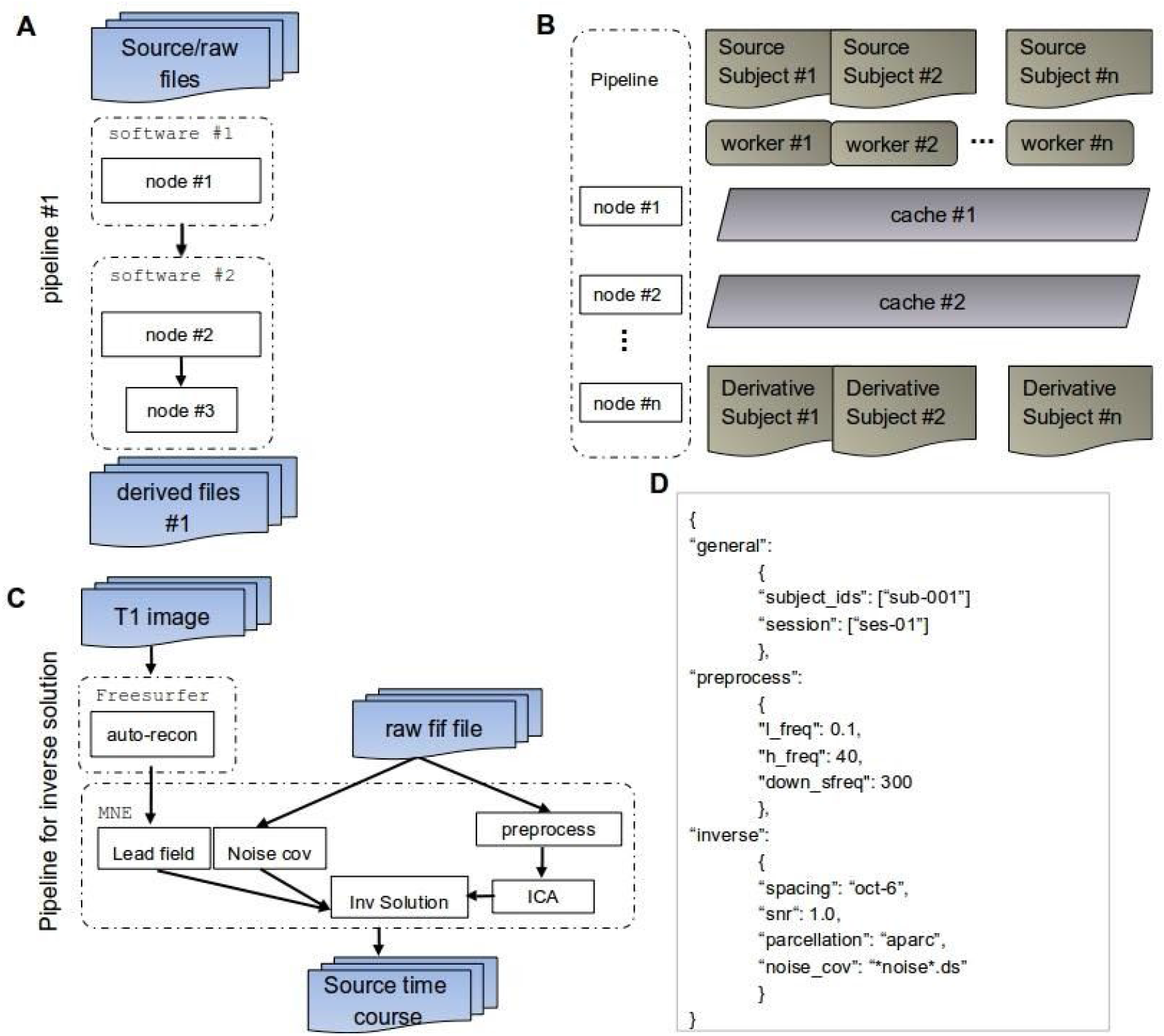
**A.** NeuroPycon offers reusable NiPype workflows for MEG/EEG processing and functional connectivity pipelines. It interfaces different software packages by linking nodes connected in acyclic graphs. The output of one pipeline can be provided as inputs to another pipeline **B.** The NiPype engine allows NeuroPycon to process subjects in parallel. If the processing interrupts due to an error, NeuroPycon will recompute only those nodes which do not have a cache. **C.** Illustrative example of ephypype pipeline for processing raw MEG data and anatomical T1 MRI to produce an inverse solution. D. Example of json file with the parameters used in the analysis. The nested structure keeps track of which pipeline corresponds to the parameters.

Although based on the same Nipype framework, graphpype and ephypype can be used separately. We designed the software with these two distinct packages so that users who only need the functionalities of one, don’t need to bother installing useless components. For instance, users who only want to make graph-theoretical analysis on fMRI data will not be obliged to install mne-python, and conversely MEG users who do not plan to do graph-theoretical analysis will not have to install radatools. Furthermore, having separate packages makes maintaining and contributing to the packages easier.

#### Software download, free license and documentation

NeuroPycon is freely available to the research community as open source code via GitHub (https://github.com/neuropycon). A basic user manual and some example scripts can be found in the tutorial webpages: https://neuropycon.github.io/ephypype/index.html, https://neuropycon.github.io/graphpype/index.html.

The documentation is built using sphinx^2^, a tool developed for python documentation that uses reStructuredText as markup language; an extension of sphinx, sphinx gallery^3^, was used to create an examples gallery by structuring the example scripts to automatically generate HTML pages.

Graphpype and Ephypype are provided under the permissive BSD 3-Clause license, which allows the use, modification and re-usability, under the condition of propagating the license. Furthermore, the BSD license allows to use the package for both commercial and non-commercial purposes.

#### Unit tests, continuous integration and python coding standards

A lot of effort has been put into providing high-quality code that is kept intact through tests and continuous integration (CI). Most of the functions and classes of the package are covered by unit-tests (as of writing this article: 68% and 70% coverage for ephypype and graphpype respectively), including tests on several kinds of data (Nifti MRI files, FIF MEG files, NumPy arrays and Radatools files). All the code also conforms to a standard Python coding convention known as PEP8 which facilitates readability and consistency with software packages in the Python scientific ecosystem.

### 2.2 NeuroPycon Packages

#### Ephypype

Ephypype is a package designed to analyze electrophysiological data using the Nipype engine. In particular, it focuses on MEG/EEG data and exploits many functions from the MNE-Python package (Gramfort et al. 2013), as well as a range of standard Python libraries such as NumPy (Van Der Walt et al., 2019) and SciPy (Virtanen et al., 2019). Current implementations allow for MEG/EEG data import, pre-processing and cleaning by automatic removal of eyes and heart related artefacts, source reconstruction, as well as sensors or source-level spectral connectivity analysis and power spectral density computation. The ephypype package features a convenient and sophisticated command-line interface which is designed to make the best use of UNIX shell capabilities and NiPype framework for parallel processing of MEG/EEG datasets. In brief, the command-line interface utilizes pattern-matching capabilities of UNIX shell to select files we want to process from the nested folder structure of a dataset and then dynamically creates a processing pipeline combining computational nodes defined in the ephypype package. In addition to being convenient, the command-line interface enables users with little programming background to easily create complex analysis pipelines that process hundreds of subjects, through a single command line.

#### Graphpype

The *graphpype* package includes pipelines for graph theoretical analysis of neuroimaging data. Computations are mostly based on radatools, a set of functions to analyze Complex Networks (http://deim.urv.cat/~sergio.gomez/radatools.php). Radatools comes as a set of freely distributed binary executables, obtained from compiling from a library originally written in Ada, and the executables are thus wrapped as command line nodes in nipype.

Although it was initially mainly used on fMRI data, the *graphpype* package can be used for the computation of graph metrics for multiple modalities (MEG, EEG, fMRI etc). This package has been developed to address the needs of functional connectivity studies that would benefit from the computation of a wide graph-theoretical metrics, including modular partitions.

It is important to keep in mind that NeuroPycon pipelines can be used in a stand-alone mode but that they can also be combined within building blocks to form a larger workflow (Fig 2), where the input of one pipeline comes from the outputs of another. As an example, the inverse solution pipeline could be used as a stand-alone pipeline to perform source localization or its output could be used as input to a new pipeline that performs all-to-all connectivity and graph analysis on the set of reconstructed time series. In principle, each pipeline, is defined by connecting different nodes to one another, with each node being either a user-defined function or a python-wrapped external routine (e.g. MNE-python modules or radatools functions).

**Figure 2.**
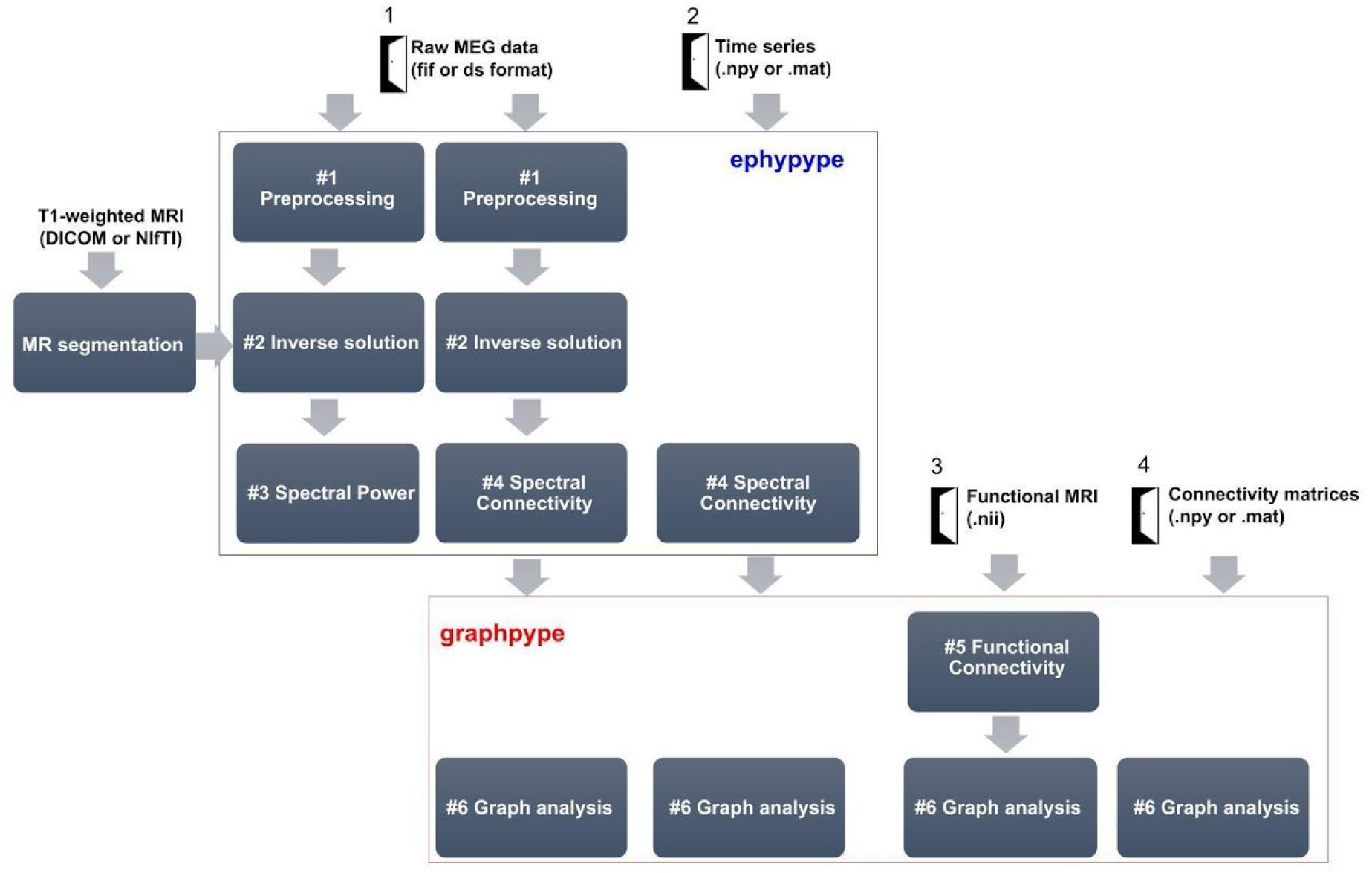
Four doorways to NeuroPycon: Illustrative pipelines showcasing different distinct uses depending on the type of input data: Full MEG preprocessing, source estimation, connectivity and graph analysis starting from raw MEG data (door 1), only connectivity and graph analysis from time series (door 2), functional connectivity and graph analysis on fMri data (door 3) or only graph analysis from connectivity matrices computed elsewhere (door 4).

### 2.3 Data import

Electrophysiology data that one would want to analyze with NeuroPycon can be in various forms and distinct data formats. NeuroPycon is designed to be able to import (1) raw MEG/EEG data, (2) time series, (3) fMRI data (4) connectivity matrices. Figure 2 showcases four different pipeline options depending on these four distinct import levels (doors). The formats required for the first option are primarily Elekta (.fif) or CTF (.ds), but import from BrainVision data (.vhdr/.eeg extension) or ascii format, is also available. New different file formats can be easily added by wrapping the corresponding import functions. Import options 2 and 4 (i.e. either time series or connectivity matrices) support.mat (Matlab) and .npy (NumPy) formats, while fMRI data import support nifti files (Fig 2, option 3). Data import for the second option (Fig 2, door 2) expects data structure of sensor or source-level time-series, allowing to read in data that may have already been analyzed by other software (e.g. BrainStorm (Tadel et al., 2011) or FieldTrip (Oostenveld et al., 2011)), and then connectivity data (possibly followed by graph metrics) are calculated using appropriate pipeline of NeuroPycon. The user can alternatively directly import connectivity matrices (Fig 2, door 3) computed elsewhere and solely use NeuroPycon to compute graph-theoretical metrics.

## 3. Presentation of the main pipelines

We now introduce the main processing pipelines that are currently proposed in the NeuroPycon software suite. These can be seen as pipeline templates or building blocks. Each pipeline may be used either independently (using its own data grabber node) or in combination. A data grabber node allows the users to define flexible search patterns, which can be parameterized with user defined inputs (such as subject ID, session, etc.). It is also possible to reassemble the preprocessing steps as required. For example, we can directly feed the pre-processed sensor-space signals to the spectral connectivity pipeline (i.e. skipping the source estimation step).

In the online documentation, we provide some key example scripting which allow the user to interactively run implementations. All example scripts are based on one of a sample MEG dataset from the OMEGA project (https://www.mcgill.ca/bic/resources/omega). The original data format follows BIDS specification (K. J. Gorgolewski et al., 2016; Niso et al., 2018). The NeuroPycon parameters settings (e.g. for the connectivity method, the source reconstruction algorithm, etc.), which are necessary for each example script, are defined in a *json* file that can simply be downloaded from the documentation page. JSON format has the advantage that it can be edited manually, and validated using external tools such as https://isonlint.com/.

### 3.1 Data preprocessing pipeline (ephypype) [Pipeline #1]

The preprocessing pipeline performs filtering, it optionally down-samples the MEG raw data and runs an ICA algorithm for automatic removal of eye and heart related artifacts. The implementation is primarily based on the MNE-Python functions that decompose the MEG/EEG signal by applying the FastICA algorithm (Hyvarinen, 1999.). An HTML report is automatically generated and can be used to correct and/or fine-tune the correction in each subject. The inclusion and exclusion of more ICA components could be performed either by re-running the same preprocessing pipeline with different option parameters (recommended option) or interactively in a Jupyter notebook (or IPython). In this last case, we suggest to save the new ICA solution and cleaned data in the corresponding node folder. Figure 3 shows the ICA decomposition obtained by running the pipeline on the sample dataset. The corresponding example script can be download as Jupyter notebook in the documentation website https://neuropycon.github.io/ephypype/auto_examples/plot_preprocessing.html

**Figure 3.**
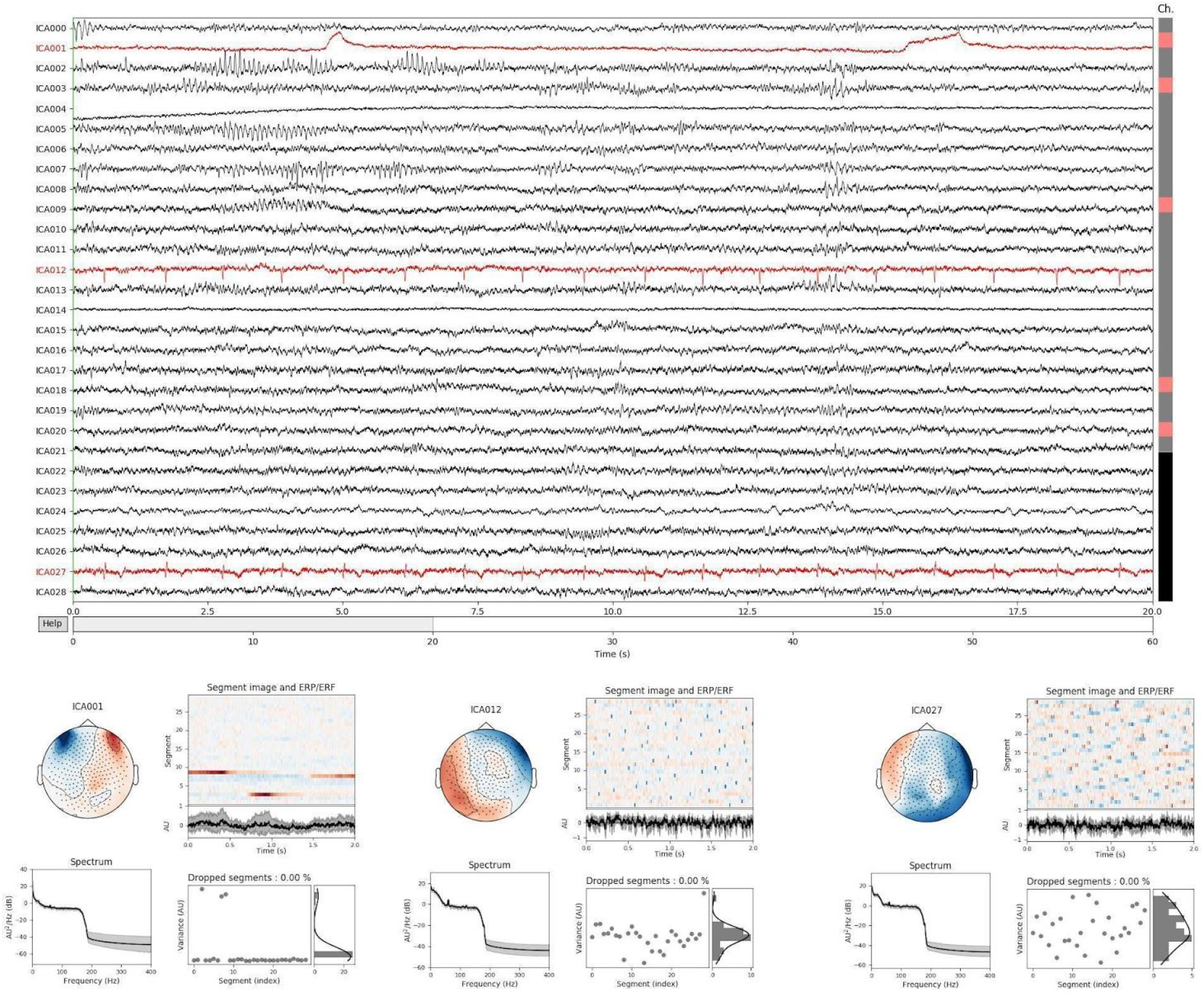
The preprocessing pipeline runs an ICA algorithm for an automatic removal of ocular and cardiac artifacts. Here we show the time series, topomap and some properties (e.g. power spectrum) of ICA components obtained by running the example script provided in the documentation website (https://neuropycon.github.io/ephypype/auto_examples/plot_preprocessing.html) on the raw MEG sample dataset.

### 3.2 Source reconstruction pipeline (ephypype) [Pipeline #2]

The inverse solution pipeline performs the source reconstruction step, i.e. the estimation of the spatio-temporal distribution of the active neural sources starting either from the raw/epoched data specified by the user, or from the output of the preprocessing pipeline (the cleaned raw data). The output of the source reconstruction pipeline will be the matrix of the estimated sources time series that could alternatively also be used as input of the spectral connectivity pipeline to study functional connectivity.

The nodes of the inverse solution pipeline wrap the MNE python functions performing the source reconstruction steps, i.e. the computation of the lead field matrix and the noise covariance matrix. These matrices are the main ingredients to solve the MEG/EEG inverse problem by one of the three inverse methods currently available in the ephypype package: MNE (Hämäläinen et al., 1993, Lin et al., 2006), dSPM (Dale et al., 2000), sLORETA (Pascual-Marqui 2002).

In particular, the lead field matrix is computed by the Boundary Element Method (BEM) (Gramfort et al., 2010) provided in MNE-Python. We use a single layer, i.e. the brain layer for MEG data, while for EEG datasets a three compartment BEM (scalp, skull and brain layers) is chosen. A graphical depiction of the source reconstruction pipeline with its node and connections is shown in Figure 4.

**Figure 4.**
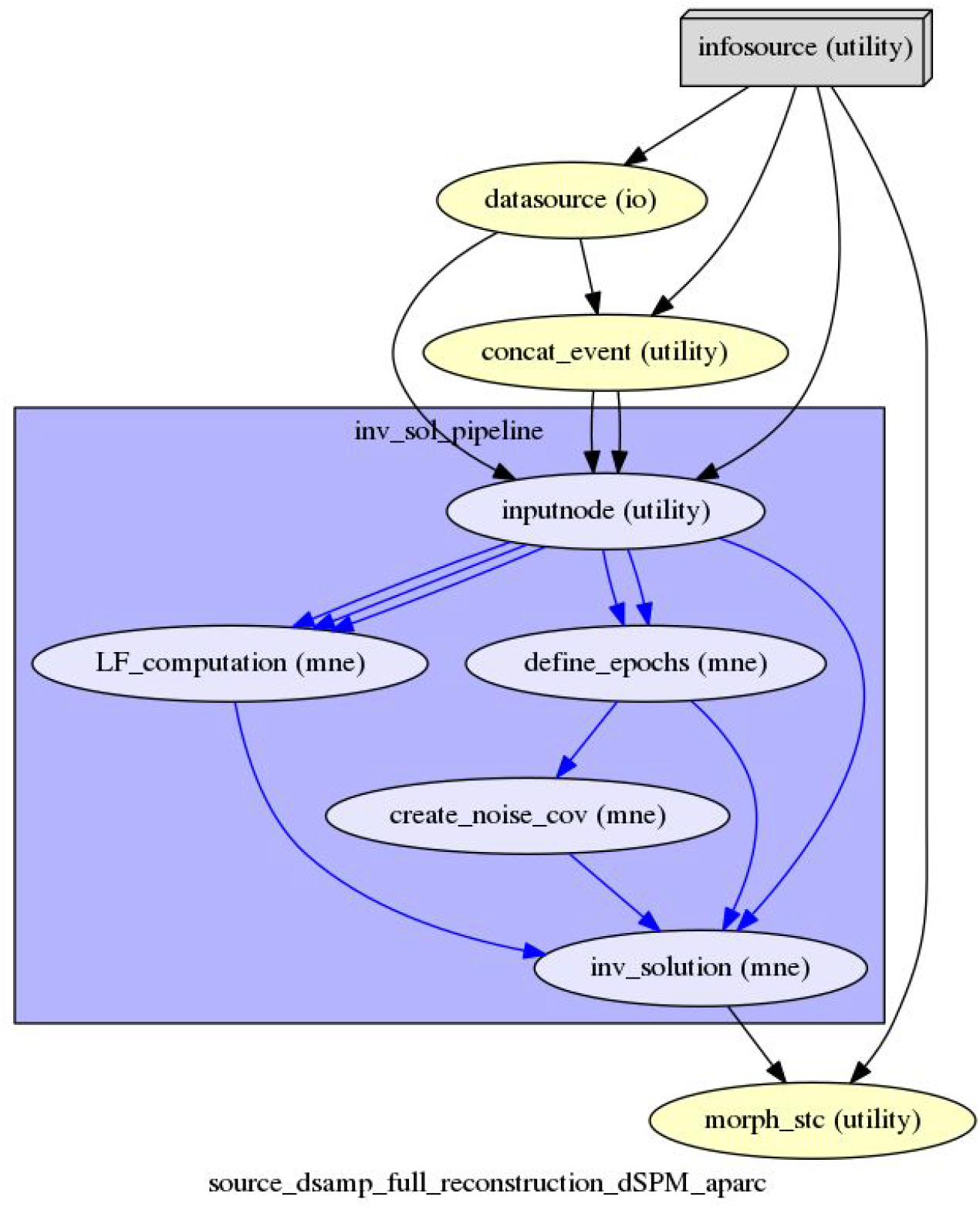
Graphical depiction of the source reconstruction pipeline with its node and connections.

To use this pipeline, a user would either need a template MRI or the individual anatomical data. In the latter case, the segmentation of the anatomical MRI has to be performed by Freesurfer in order to generate surfaces and parcellations of the structural data. The anatomical segmented data will be used in the pipeline to extract the BEM surfaces and to create the source space. By default it is expected that the current dipoles are situated on the cortical surface, but it is also possible to set a mixed source space constituted by the cortical surface and the volumes of some user-selected subcortical regions, as amygdala, thalamus, cerebellum, etc. Finally, the segmented MRI data are also used to perform the coregistration step between the MEG/EEG and MRI coordinate system. This is the only manual step one has to perform before using the source reconstruction pipeline and can be performed by *mne_analyze* process of MNE-C^4^ or an MNE-Python Graphical User Interface (GUI). In the future, automatic co-registration will become available when this feature becomes a robust function in MNE python. A template MRI, just like the one provided by the Freesurfer software, could be used if the individual anatomical data are not available. Figure 7 shows the results obtained by running the inverse solution pipeline on the sample dataset. The script used to generate this figure can be downloaded as Jupyter notebook from the documentation website https://neuropycon.github.io/ephypype/auto_examples/plot_inverse.html.

### 3.3 Spectral power pipeline (ephypype) [Pipeline #3]

The power pipeline computes the power spectral density (PSD) in either sensor or source space. It also computes the mean PSD for each frequency band specified by the user. The latter can choose to compute the PSD by Welch’s method or multitapers. The input of the pipeline can be either raw/epoched data specified by the user or simply the output of another pipeline, e.g. the cleaned raw data from the preprocessing pipeline or the estimated source time series from the source reconstruction pipeline. Figure 6. A shows the results obtained by running the power pipeline on the sample dataset. The script used to generate this figure can be downloaded as Jupyter notebook in the documentation website https://neuropycon.github.io/ephypype/autoexamples/plotpower.html.

### 3.4 Spectral connectivity pipeline (ephypype) [Pipeline #4]

The spectral connectivity pipeline computes the connectivity matrices in the frequency domain. The current implementation is based on the spectral connectivity computation in MNE, and can be computed on time series in NumPy format (in either sensor or source space), or even from Matlab format after conversion using the SciPy package. All the frequency-domain coupling measures available in MNE-Python are directly accessible through this pipeline (e.g. Coherence, Imaginary Coherence, Phase Locking Value, Phase-Lag Index). Figure 6.B shows the results obtained by running the connectivity pipeline on the sample dataset. The script used to generate this figure can be downloaded as Jupyter notebook in the documentation website https://neuropycon.github.io/ephypype/auto_examples/plot_connectivity.html.

### 3.5 Functional connectivity analysis (graphpype) [Pipeline #5]

Starting from pre-processed functional MRI data, we provide a pipeline named “nii_to_conmat”, which allows the user to compute functional connectivity from a given file in Nifti format, and a corresponding template. The pipeline convolves the template with a gray-matter mask, extracts time series under the restriction of sufficient proportion of voxels in the template ROI, regresses non-interest covariates (white matter, cerebrospinal fluid, movement parameters, etc) and computes Z-score Pearson correlations over residuals signals. An example of this pipeline, followed by the graph-theoretical analyses pipeline (see following section) is provided at https://neuropycon.github.io/graphpype/auto_examples/plot_nii_to_graph.html#nii-to-graph

### 3.6 Graph-theoretical analyses pipeline (graphpype) [Pipeline #6]

Once a connectivity matrix has been obtained, the most common step to compute graph theoretical (GT) analyses involves a form of thresholding to “sparsify” the matrix and keep only the most relevant edges. One classical thresholding option is the use of a density-based thresholding (e.g. only keep edges with the 5% highest connectivity values)(Rubinov and Sporns, 2010, Bassett and Lynall, 2013). Another possibility is fixed value thresholding (e.g. all coherence values lower than 0.5 are put to zero). For signed metrics, such as Pearson correlation, the sign will be taken into account at this step.

The computation of standard GT metrics (includes weighted and signed versions) primarily relies on wrapping one of the most efficient modularity optimisation software tools called Radatools (http://deim.urv.cat/~sergio.gomez/radatools.php). Radatools offers the possibility to compute GT properties for several classes of networks (binary/weighted, unsigned/signed) and allows the computation of most global (e.g. mean path length, global efficiency, clustering coefficient, assortativity, as well as their weighted counterparts) and nodal metrics (e.g. degree, betweenness centrality, etc.). Radatools is mostly known for highly efficient modularity optimisation possibilities. In addition to being amongst the few tools to offer modular partition on weighted signed networks, it also offers the choice of several algorithms ranging from lower quality with a fast execution algorithm (Newman et al, 2006) to exhaustive searches (high quality but time consuming). Interestingly, it is possible to define a sequence of these algorithms to combine the advantages of these algorithms. Some specific metric computation, mostly related to node roles (participation coefficient and within-module normalized degree, see Guimera et al, 2005) have also been specifically coded and are part of Radatools’ standard modular decomposition pipeline. The script used to generate this figure can be downloaded as Jupyter notebook on https://neuropycon.github.io/graphpype/auto_examples/plot_invts_to_graph.html.

Another package allowing for graph-theoretical analysis, very popular in the neuroscience community, is the Brain Connectivity Toolbox (Rubinov and Sporns, 2011). As an example of a wrap, and to showpotential users what would be done to integrate their favorite software functions, we documented the definition of a small pipeline consisting of one node (K-core from the bctpy package): https://neuropycon.github.io/graphpype/how_to_wrap.html

## 4. Validation of NeuroPycon group analysis using previously published work on open data

In order to show how to use NeuroPycon pipelines to analyze a cohort of subjects, we replicated the results obtained by Jas et al., 2018. In this article, the authors reanalysed an open dataset from Wakeman and Henson (2015) using the MNE software package, with the aim of providing group analysis pipelines with publicly available code and documentation. The data consist of simultaneous MEG/EEG recordings from 19 healthy participants performing a visual recognition task. Subjects were presented images of famous, unfamiliar and scrambled faces. Each subject participated in 6 runs, each 7.5 min in duration. The data were acquired with an Elekta Neuromag Vectorview 306 system. We focused only on MEG data and used NeuroPycon pipeline #1 to preprocess the MEG raw data and pipeline #2 to perform source reconstruction of time-locked event-related fields.

In the following, we describe the main steps of the analysis. All parameters of the analysis are set in a single json file, with a nested structure to keep track of which parameters correspond to which steps. All the scripts can be downloaded from https://github.com/neuropycon/neuropycon_demo.git. Note that anatomical segmentation is performed using a dedicated script and the preprocessing and source reconstruction pipelines are combined in a single workflow. The individual steps of this group-analysis replication with NeuroPycon are described below.

### 4.1 Cortical segmentation

The solution of MEG inverse problem requires knowledge of the so-called lead field matrix. A cortical segmentation of the anatomical MRI is necessary to generate the source space, where the neural activity will be estimated. A Boundary Element Model (BEM) which uses the segmented surfaces is used to construct the lead field matrix. To perform the cortical segmentation we provide a workflow based on nipype wrapping the recon-all command of Freesurfer. The output of recon-all node is used as input of another node that creates the BEM surfaces using the FreeSurfer watershed algorithm (Segonne et al., 2004). The workflow generates an HTML report displaying the BEM surfaces as colored contours overlaid on the T1 MRI images to verify that the surfaces do not intersect.

The main advantage to use this workflow lies in the parallel processing provided by nipype engine, that allows segmenting the 19 MRI data in less than two days while processing a single MRI generally takes one day.

### 4.2 MEG data processing and Independent Component Analysis (ICA)

The data provided by OpenfMRI were already processed using the proprietary Elekta software MaxFilter (Taulu, 2006), used to remove environmental artifacts and compensate for head movements. We used these data as input of our preprocessing pipeline (pipeline #1). Since we want to study event-related fields, a bandpass filter was applied to the data between 0.1 to 40 Hz, without downsampling. The pipeline #1 also runs an ICA decomposition on filtered data to remove cardiac and ocular artifacts. The names of EoG and ECG channels, the number of ICA components specified as a fraction of explained variance (0.999) and a reject dictionary to exclude time segments were set in the json parameter file. While in Jas et al. (2018) the ICA solution is used to directly remove the bad components from epoched data, we perform this operation on the unsegmented data. However, due to the linearity of ICA operation, this does not affect the results.

### 4.3 Extracting events

For the sake of reproducibility, following Jas et al. (2018) we use an auxiliary NeuroPycon node (concat_event, see Figure 4) to (i) extract the events from the stimulus channel ‘STI101’ and (ii) concatenate the six different runs for each subject. The output of this node is the input of the source reconstruction pipeline described below.

### 4.4 Source estimate

Figure 4 shows the graph corresponding to the source reconstruction pipeline (pipeline #2). Before running this pipeline, the coregistration between the MRI and MEG device needs to be performed. As highlighted in the section describing the source reconstruction pipeline, this is the only manual step before using the pipeline and it represents a critical step to obtain a good localization accuracy. Similarly to Jas et al. (2018), we used the coregistration file provided by Wakeman and Henson (2015). The coregistration file is used in the LF_computation node where the source space and eventually the BEM surfaces are also created. As source space, we choose a dipole grid located in the cortical mantle. By setting in the json file ‘oct-6’ for the ‘spacing’ parameter leads to around 8196 vertices in the source space for each subject. Since we are analyzing MEG data a single layer head model with only the inner skull surface is sufficient for the BEM computation. The noise covariance matrix is estimated from 200 ms of prestimulus data. Since we want to do source estimation in three different conditions (famous faces, unfamiliar faces and scrambled), we provided all information related to the events in the json file. We also specified as inverse method dSPM that was one of the inverse methods used by Jas and colleagues. Finally, in the morph_stc node (Figure 4) the output of the inverse solution pipeline, i.e. the reconstructed neural time series is morphed to the standard FreeSurfer average subject named fsaverage.

### 4.5 Comparison

Figure 5 (right) shows group average of dSPM solutions for the contrast between both types of faces together and scrambled at 170ms post-stimulus. The image was produced by subtracting normalized solutions of faces to the ones of scrambled. The results are similar to the ones obtained by Jas and colleagues (Figure 5, left).

**Figure 5.**
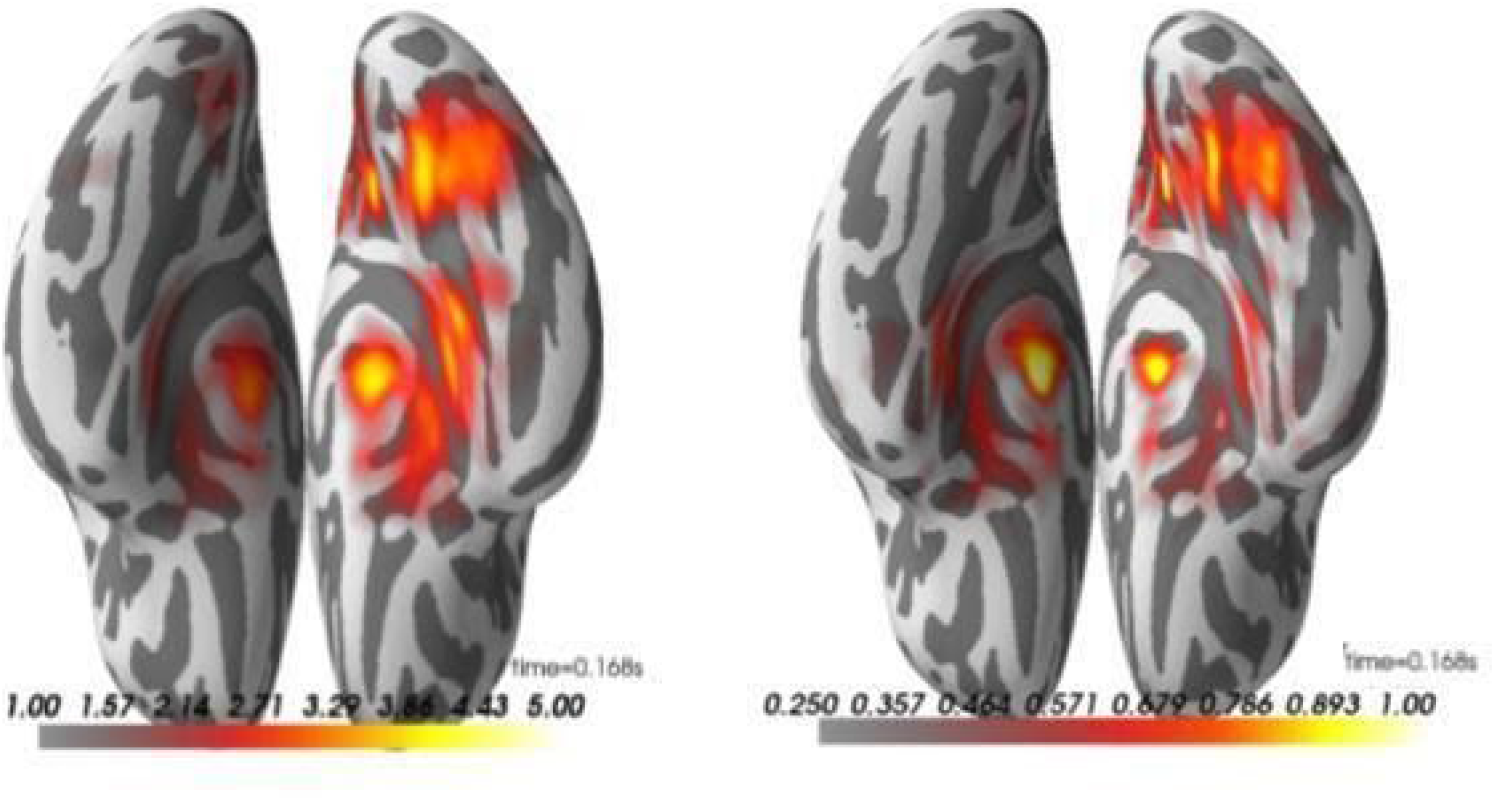
Group average on source reconstruction with dSPM obtained by Jas et al. 2018 (Left) and using NeuroPycon pipelines (Right). (see analysis implementation and details in Section 4)

We also measured the computational time needed to compute ICA by MNE and NeuroPycon with the aim to see the performance benchmarks of running MNE python in NeuroPycon versus in pure python. While the parallel library used by Jas and colleagues leads to a computational time comparable with the one obtained by using NeuroPycon, one of the main advantages provided by NeuroPycon is related to the caching provided by Nipype engine that stores intermediate files and recomputes only those nodes which do not have a cache. This has a significant impact on the speed of the analyses mainly if some error occurs in the analysis of some subjects. Furthermore, another advantage of Nipype engine is the ability to easily switch from MultiProc (i.e. multiproc computer) to queuing cluster systems such as SGE or SLURM.

## 5. Postprocessing connectivity and graph-theoretical metrics

### 5.1 Gathering results

The output of a NeuroPycon pipeline results in a specific directory architecture, where all the results of each iteration are sorted by nodes. A post-processing step allows us to gather results in a handy way for subsequent statistical analyses and further result representation and visualization. For the graph-metrics computed via graphpype, the most straightforward way is to gather graph-based results in a dataframe, for further processing outside in Excel (TM) or R. The postprocessing tools allowing for generic post-processing steps can be found in the “gather” directory of ephypype and graphpype.

### 5.2 Statistical analyses

Within NeuroPycon, it is possible to conduct group-level statistics in several ways. Any current or future statistical analysis functions available through tools wrapped by NeuroPycon are automatically available for use in NeuroPycon pipelines (e.g. all statistical analyses provided by MNE python). Additionally, the graphpype package offers several functions to compute statistics on connectivity and graph data. These include assessing statistical significance by computing parametric tests (paired and unpaired t-test, binomial, Mann-Whitney, etc.) between groups of matrices or vectors. Of course, with the increasing number of dimensions we also need to address the multiple comparison problem. It is possible to compute several levels of significance accounting for multiple comparisons, tailored for connectivity and graph metrics: For instance, a False Positive metric (1/#of tests) has been suggested to be an acceptable threshold when hundreds or even thousands of nodes are at play (Bassett and Lynall, 2013). Other implemented tools include False Discovery Rate (Benjamini and Hochberg) and Bonferroni correction. An alternative approach is of course to implement non-parametric permutation testing over mean connectivity matrices. The time-consuming steps of permutation computing herecritically benefit from the parallelization available in NeuroPycon (via Nipype engine). A reasonable number of computations (e.g. 1000) can be achieved in a relatively short time (a few hours for the full network pipeline computation, including modular decomposition, assuming typical network sizes of ∼100 nodes). The gather_permuts module of graphpype package in the “gather” directories offer a range of functions allowing for computation of the corresponding p-values.

### 5.3 Visualization tools (graphpype, visbrain)

Numerous tools can be used to visualize the results and data computed by or manipulated within NeuroPycon. One option is to use visualization tools currently used within MNE python, such as pysurfer^5^ and mayavi (Ramachandran and Varoquaux, 2011), as well as a more recent implementation using PyVista (Sullivan & Kaszynski, 2019). An example of visualization provided by MNE python is shown in Figure 3 containing the output of preprocessing pipeline, i.e. the topographies and time series of the ICA components. Another more recent option to visualize the results is to use visbrain (http://visbrain.org), a python based open-source software dedicated to the visualization of neuroscientific data (Combrisson et al., 2019). It is built on top of PyQt^6^ and VisPy (Campagnola et al., 2015), a high-performance visualization library that leverages the Graphics Processing Units (GPU). Below we describe visualization procedures for different data generated with NeuroPycon using visbrain (pysurfer and mayavi descriptions can be found elsewhere).

The functions of Visbrain are not wrapped as nodes, as Visbrain is conceptually used as post-processing steps outside of NeuroPycon. In the future, Visbrain capabilities may be used in standardized reports if the need arises.

#### 5.3.1 Visualization of sensor-space data

Figure 6 uses the source object (SourceObj) class of visbrain that allows to represent MEG sensors and assign additional data values to each one of them. One available option is to represent the input data (i) through a color bar and (ii) a marker radius proportional to its amplitude. Figure 6.A depicts the output of the spectral power pipeline (see section 3.3), where the PSD was computed in sensor-space over three different frequency bands (theta, alpha and beta). Here we show the results on alpha band. By contrast, Figure 6.B is generated using the source object class together with the visbrain’s connectivity object (ConnectObj) used to draw connectivity lines between nodes. Here we show the results of the connectivity pipeline, i.e. the sensor-level connectivity matrix obtained by computing the coherence among MEG sensors in alpha band.

**Figure 6.**
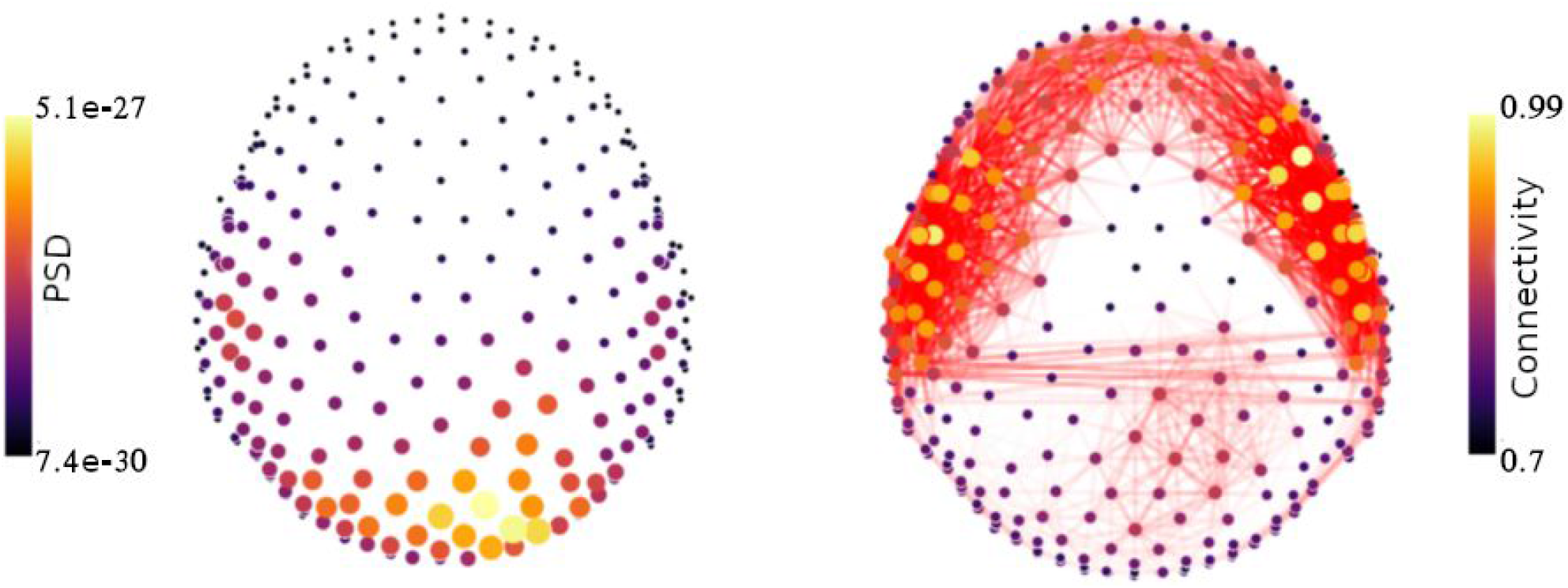
**(A)** The spectral power pipeline computes single-trial and mean PSD for each selected frequency band. Here we show illustrative results of computing alpha power running the NeuroPycon power pipeline template on the sample MEG data. The size and color of each sensor vary with the alpha power value. The script used to generate this figure is provided in the documentation website: https://neuropycon.github.io/ephypype/auto_examples/plot_power.html. **(B)** The connectivity pipeline performs connectivity analysis in sensor or source space. Here we show illustrative results obtained using the connectivity pipeline to compute coherence between MEG sensors in alpha band. Connectivity edges are colored according to the strength of the connection, while the node size and color depend on the number of connections per node. These results are obtained by running the examples script in the documentation webpage: (https://neuropycon.github.io/ephypype/auto_examples/plot_connectivity.html,

#### 5.3.2 Visualization of source space data

To achieve 3D brain visualizations, the output data resulting from the different pipelines (power, connectivity, inverse solution and graph) can be interfaced with the *Brain* module of Visbrain (http://visbrain.org/brain.html) that can be used to visualize the estimated source activity, connectivity results, PSD on source space, and graph analysis results. Illustrative figures that can be produced by the *Brain* module are shown in Figures 7 and 8. Figure 7 shows the output of the inverse solution pipeline, i.e. the reconstructed neural activity in each ROI of a user-defined atlas (Desikan-Killiany Atlas) at a given time point. Figure 8 shows a graph obtained for alpha band after computing functional connectivity between all pairs of regions starting from the ROI estimated time series computed by the inverse solution pipeline. The graph is obtained after thresholding at 5% highest coherence values.

**Figure 7:**
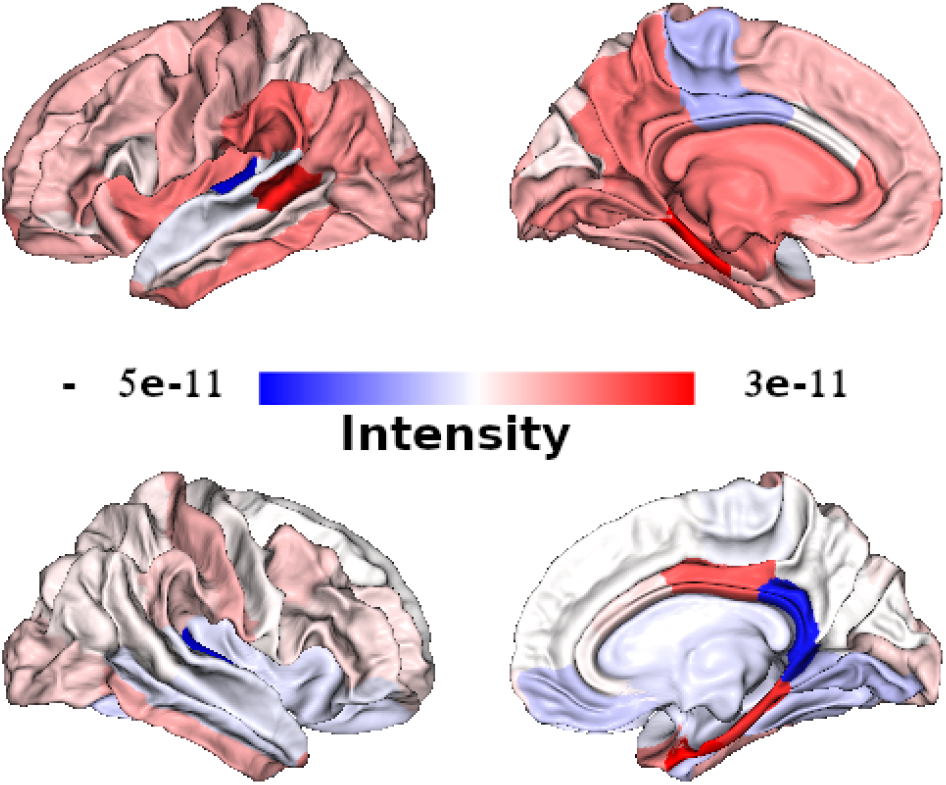
Here we show the output of the inverse solution pipeline, i.e. the reconstructed neural activity at a given time point, in ROIs from a user defined atlas (Desikan-Killiany Atlas). The results are obtained by running the example script in the documentation webpage (https://neuropycon.github.io/ephypype/auto_examples/plot_inverse.html).

**Figure 8:**
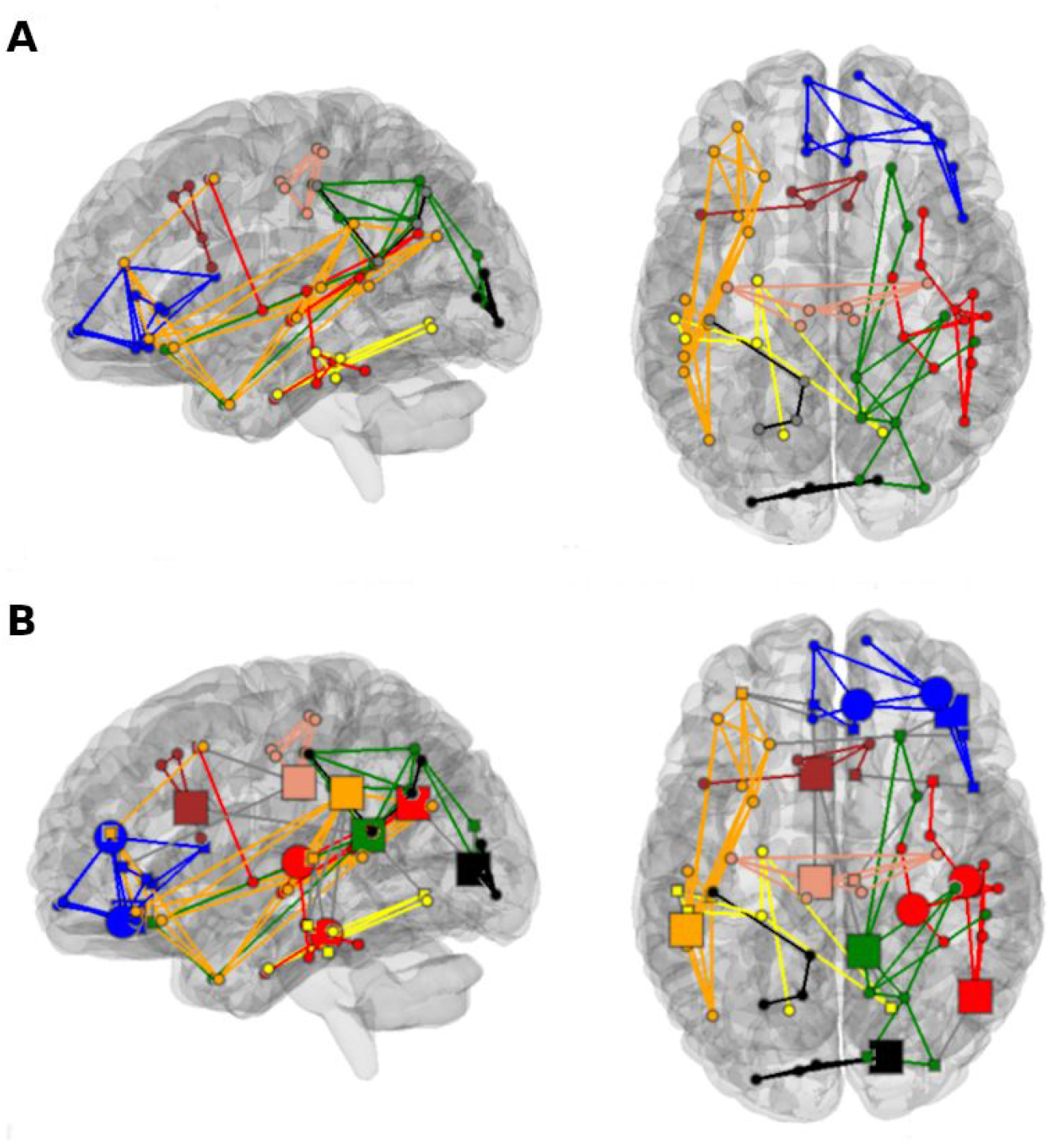
Representation of a graph obtained from resting-state MEG data for alpha band (left part = seen from left; right part = seen from top) after computing functional connectivity between all pairs of regions (Figure 7). The graph is obtained by retaining the 5% highest coherence values. The results of modular decomposition are displayed with the same color for the edges between two nodes in the same module, and in grey for edges between nodes belonging to different modules. Two representations of the same results are displayed: with modules (panel A), and with modules and node roles definition of Meunier *et al.* (2009) as the shape (square = connector) and size (bigger shape = hub) of the nodes (panel B). In the lower part, inter-modular edges are represented in grey. From a given size in decreasing order, modules are all represented in black.

## 6. NeuroPycon Command Line Interface (CLI)

As previously mentioned, the construction of neuropycon data processing pipelines is done in a python script specifying the affinity and arguments of the processing nodes and the source of input data. Such scripts can be distributed, shared and reused later, which facilitates reproducibility and results sharing. However, this can arguably sometimes be tedious if we want to construct and run simple pipelines quickly. To address this problem, we’ve provided neuropycon with a Command Line Interface (CLI) which is provided in the ephypype package. Indeed, at the moment the command line interface wraps only some of the functionality of the ephypype package only but will be expanded in the future.

CLI is aimed at building the processing workflows on the fly leveraging the UNIX shell wildcards mechanism for flexible input specification (Fig. 8). It wraps the processing nodes of ephypype together with their options and arguments exposing to the end user a subcommand for each node. Specifying these subcommands in order, the user in effect chooses the desired processing steps which are assembled together into the nipype workflow at the command invocation. More precisely, we provide the terminal command *neuropycon* which is followed by the sequence of subcommands corresponding to the desired processing nodes with specified options and arguments for each. This chain of subcommands to *neuropycon* determines the specific form of the processing pipeline we want to apply to our data.

In practice, the use of the neuropycon terminal command looks like the example shown in Figure 9, where a sequence of commands segments the data into 1 second epochs, converts them to Numpy format and computes the default connectivity measure in the 8-12 Hz frequency band.

**Figure 9:**
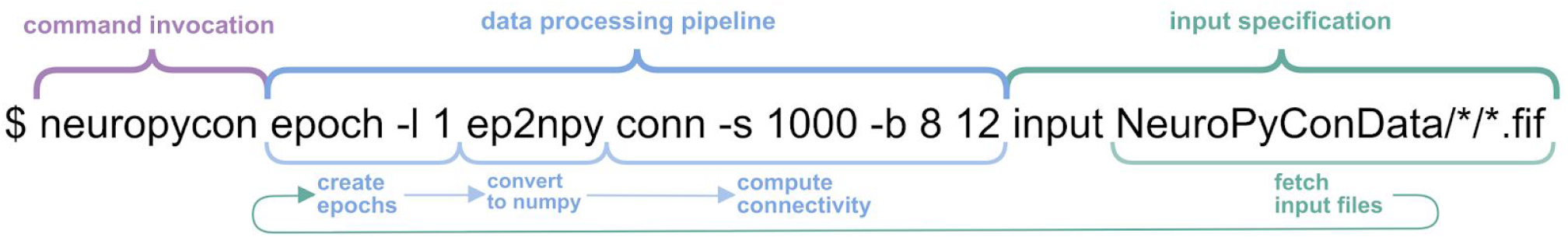
Example of the CLI command computing connectivity metrics on a group of files. This command grabs all the .fif files in the two-level nested folder structure, creates one-second epochs from them, converts the epochs to NumPy arrays format, performs a default connectivity metrics calculation -between 8-12 Hz-on the converted data and saves the results.

The command can be split into three functional blocks. First goes the command name *neuropycon.* Then it is followed by a chain of processing subcommand for each of which we specify options and arguments unique to each processing node. In the example depicted in Figure 9, the supplied processing subcommands are *epoch, ep2npy* and *conn,* which perform data epoching, conversion to NumPy format and spectral connectivity computations. The last block is always the input specification. Although the input node really goes first in the stream of data processing, putting the input specification to the rightmost position of the composite command allows us to specify an arbitrary number of input files to the pipeline which is beneficial when working with wildcards matching.

The input block always starts with the *input* subcommand and is followed by a list of file paths we are applying the processing pipeline to. In the example shown in Figure 9, the list of files is specified using the UNIX wildcards matching mechanism and can be spelled out as ‘Go to each subfolders of the NeuroPyConData folder and take all the ‘.fif’ files contained in it’ (here we presume that there’s only one level of nesting in NeuroPyConData folders structure, i.e. files are organized according to the following scheme: NeuroPyConData/<SubjectName>/<subject_data.fif>).

Integration with the UNIX wildcards pattern matching is one of the biggest strengths of the supplied CLI since it allows for flexible and concise fetching of files in the nested folders hierarchy given that these folders are organized in a regular and well-defined fashion, which is often the case for electrophysiological datasets. A more detailed explanation of the command line interface operation principles and examples can be found on the documentation webpage (https://neuropycon.github.io/ephypype/cli.html#neuropycon-cli).

## 7. Strengths of NeuroPycon and advantages of its Nipype-based framework

NeuroPycon is based on the Nipype engine and fully adheres to its architecture and global software philosophy. In this section, we will here briefly summarize the rationale and key components of Nipype, and then outline the strengths and added-value that NeuroPycon brings to the community through this architecture.

### 7.1 Nipype in a nut-shell

Nipype is an open-source, community driven, python-based software package that enables interactions between existing neuroimaging software in a common framework and uniform semantics (Gorgolewski, et al, 2015). The design of workflows using Nipype allows for intuitive and tractable implementations of even quite complex processing pipelines. To appreciate concrete advantages that NiPype confers to NeuroPycon, it is useful to briefly overview Nipype’s three main components: **(I)** *Interfaces* to external tools that provide a unified way for setting inputs, executing and retrieving outputs. The goal of Interfaces is to provide a uniform mechanism for accessing analysis tool from neuroimaging software packages (e.g. Freesurfer, FSL, SPM, etc). **(II)** A *workflow engine* allows to create analysis pipelines by connecting inputs and outputs of interfaces as a directed acyclic graph (DAG). In order to be used in a workflow the Interfaces have to be encapsulated in node objects that execute the underlying Interface in their own uniquely named directories, thus providing a mechanism to isolate and track the outputs resulting from the Interface execution. Nodes can be connected together within a workflow: by connecting the outputs of some node to input of another one, the user implicitly specifies dependencies. Furthermore, workflow can itself be a node of the workflow graph. Nodes provides also an easy way to implement function defined by the user. **(III)** A *plug-in* executes a workflow either locally or in a distributed processing environment. No changes are needed to the workflow to switch between these execution modes. The user simply calls the workflow run function with a different plug-in and its arguments.

### 7.2 NeuroPycon’s main assets and advantages

#### (a) Multiprocessing

the implementation of multiprocessing is very easy, and can be either made for multi-processing on a same machine with multiple cores (Multiproc plugin) or a cluster with multiple machines in parallel (q-sub/ipython plugin). In addition to substantially speeding up the computations for a planned analysis, the ability to easily launch multiprocessing computations also encourages users to rapidly test and compare different analyses options (e.g. various preprocessing strategies, or different source localisation methods, or even optimizing parameter selection, such as matrix thresholding or graph construction, or frequency band choices, etc.). Easy definition of multiprocessing is also an advantage over Matlab-based scripting when it comes to handling big datasets. While multiprocessing exists in Matlab its implementation requires specific coding, whereas in NeuroPycon the exact same code is used for both sequential and parallel processing, except for one line that specifies the option.

#### (b) Caching

Thanks to the use of Nipype, NeuroPycon stores intermediate files, and tests if the source code of each node has been modified. Hence, if a part of the pipeline is modified, only the modified parts will be recomputed. This has a significant impact on the speed of the analyses.

#### (c) Report

Each node of the workflow creates a subfolder (under the workflow directory) called _report containing a text file (report,rst) with all relevant node information, i.e. the name of the node, the input and output parameters, the computational time to execute the node.

#### (d) Choices and interaction between multiple software tools

It is not uncommon in the literature to see that graph analysis of EEG or MEG data is achieved by combining distinct independent software tools (e.g exporting connectivity data from one of the MEG/EEG analysis toolboxes available to a functional connectivity software such as the Brain Connectivity Toolbox (BCT) (Rubinov and Sporns, 2010)). By contrast, NeuroPycon provides a unified framework with seamless interactions between tools allowing to compute graph properties starting from raw MEG/EEG data.

#### (e) Expandability

It should be noted that the general « wrapping » concept makes it possible to expand NeuroPycon’s workflows to include software combinations other than those currently proposed. Extending the currently available functionalities to wrap other software packages is in theory reasonably straightforward. The biggest challenge is to ensure the compatibility of format between the packages and to code the corresponding converters. To illustrate the steps associated with wrapping new functions from other packages into NeuroPycon, we provide (i) a tutorial (see the graphype documentation) on how to wrap a single function (Kcore computation of the Brain Connectivity Toolbox), which is a metric available in BCT but not in Radatools, and integrate it in a simple pipeline: https://neuropycon.github.io/graphpype/how_to_wrap.html#how-to-wrap and (ii) an example on how to create a workflow connecting a Node that wraps a function of a Matlab toolbox(e.g. FieldTrip) with a Node that wraps a function of a python toolbox (e.g. MNE): https://neuropycon.github.io/ephypype/auto_examples/run_fieldtrip_wf.html.

On the other hand, the compatibility is readily guaranteed if the aim is to include in NeuroPycon a new pipeline based on an algorithm already implemented in a software wrapped by NeuroPycon (e.g. MNE python or radatools). For example, adding a source reconstruction pipeline using Linear Constrained Minimum Variance beamformer (Van Veen et al., 1997) as inverse method is possible via a small modification of the ephypype package (not more than ten lines of code). This type of extension is also illustrated in NeuroPycon’s online documentation, see for example: https://neuropycon.github.io/ephypype/tutorial/lcmv.html

#### (f) Multimodal analysis

NeuroPycon also provide an advantageous framework for multimodal analyses (e.g. combining electrophysiological and neuroimaging data). Indeed, in addition to its own pipelines, NeuroPycon can benefit from the interfaces already made available for neuroimaging analysis via Nipype. For example, since the latter wraps most of the functions available in Freesurfer (Dale et al., 1999), MRI segmentation and parcellation, and subsequent MEG source space processing can all be completed with a single reproducible, light-weight and shareable NeuroPycon pipeline.

#### (g) Open-source, readability and reproducibility

Because it is written in python, the NeuroPycon code is compact and highly readable. In addition, python is free (which is not the case of other high-level scientific languages such as Matlab). As a result, NeuroPycon is a freely accessible tool that can be readily used by students and researchers across the globe without the need to purchase commercial software. Being open source, NeuroPycon promotes pipeline sharing and enhances reproducibility in an open-science mind set.

## 8. Relationship to other toolboxes

NeuroPycon is not an alternative toolbox designed to replace or compete with existing software. On the contrary, the strength of NeuroPycon is that it builds upon existing tools and brings them into a unifying framework, through the Nipype engine. This has several immediate advantages over individual toolboxes: (1) NeuroPycon can be used to compare results of using different algorithms from different existing toolboxes (e.g. wrapping MNE, brainstorm, SPM and fieldtrip, into a single NeuroPycon pipeline allows for simple and direct comparison of source estimation -implemented in distinct toolboxes-in a common framework), (2) NeuroPycon pipelines can be very exhaustive, including all processing steps needed to go from raw data (functional and structural) to group-level statistics of source-space connectivity and graph-theoretical analyses. Because each pipeline is associated with a shareable parameter file with all the parameters for all the nodes, the whole analysis can easily be replicated. (3) The unifying framework of NeuroPycon also provides an ideal environment for multimodal fusion. For example, the graph package can be used with fMRI and MEG or EEG data from the same individuals and provides a practical framework to compare or integrate data from different brain imaging modalities, (4) A further advantage of NeuroPycon, compared to other available individual tools, is that it benefits from Nipype’s efficient computing functionalities, such as embedded multi-threading for parallel processing and caching, which is very useful when it comes to data sets of large cohorts, (5) In addition to benefiting from the known advantages of Python, the fact that all the analyses happen in a single programming environment means that the user does not need to be familiar with different languages that the tools are programmed in, and (6) Finally, because NeuroPycon integrates available open tools, all improvements and new functionalities thatare added to these toolboxes, will be readily available to NeuroPycon and so NeuroPycon’s strength will continually grow as the individual toolboxes continue to improve.

In sum, rather than being a competitor to existing software, NeuroPycon allows the community to benefit from the strengths of existing tools in a common open and computationally efficient framework designed to facilitate method comparison, sharing and replication of results.

## 9. Discussion

NeuroPycon is an open-source analysis kit which provides python pipelines for advanced multiprocessing of multi-modal brain data. It consists of two primary, complementary components: ephypype, which facilitates preprocessing and source localization pipelines; and graphpype, which integrates spectral and functional connectivity pipelines, as well graph analysis pipelines.NeuroPycon is based on the Nipype engine and inherits thereby its philosophy of wrapping multiple established processing software tools into a common data analysis framework. The use of NeuroPycon allows for portability, simplified code exchange between researchers, and reproducibility of the results by sharing analysis scripts.

Additionally, conducting graph-theoretical analysis in NeuroPycon allows for comparing and merging graph results from different imaging modalities and different toolboxes (which might be implemented in different programming languages).

NeuroPycon is designed for users with a reasonable knowledge of the software tools that it can wrap. but who would like to benefit from automatic implementation (ready-to-use pipelines) of features that would be otherwise more complex (or impossible) to implement. Parallel processing and caching are features that are particularly powerful and convenient when it comes to large data sets. NeuroPycon could also become very valuable for users who want to process multimodal data (e.g. MEG and fMRI) in a unified framework. Last but not least, students and researchers will hopefully find NeuroPycon to be a convenient framework to easily share MEG/EEG analysis pipelines.

In terms of ongoing and future development, we plan-among other things-to make NeuroPycon BIDS-compatible so that the inputs are BIDS datasets and the intermediate outputs comply with the upcoming BIDS derivative specification. Also, because the Nipype engine provides an important backbone to NeuroPycon, we hope to work closely with the Nipype community in the future, especially as NeuroPycon evolves and expands its functionalities. One of the current shortcomings is the limited packages that have so far been wrapped. Hopefully, the community of NeuroPycon users and developers will continue to expand, and thus increase its functionalities and range of pipelines available.

## Acknowledgements

AP was supported by a CNR short-term mobility grant. KJ was supported by funding from the Canada Research Chairs program and a Discovery Grant (RGPIN-2015-04854) from the Natural Sciences and Engineering Research Council of Canada, a New Investigators Award from the Fonds de Recherche du Québec - Nature et Technologies (2018-NC-206005) and an IVADO-Apogée fundamental research project grant. This research is also supported in part by the FRQNT Strategic Clusters Program (2020-RS4-265502 - Centre UNIQUE - Union Neurosciences & Artificial Intelligence - Quebec).

## Footnotes

1 http://deim.urv.cat/~sergio.gomez/radatools.php

2 http://www.sphinx-doc.org/en/master/

3 https://sphinx-gallery.github.io/index.html

4 https://www.mne-cpp.org/

5 http://pysurfer.github.io/

6 https://wiki.python.org/moin/PyQt

